# Olfactory and neuromodulatory signals reverse visual object avoidance to approach in Drosophila

**DOI:** 10.1101/472605

**Authors:** Karen Y. Cheng, Mark A. Frye

**Affiliations:** UCLA Department of Integrative Biology and Physiology

## Abstract

Innate behavioral reactions to sensory stimuli may be subject to modulation by contextual conditions including signals from other modalities. Whereas sensory processing by individual modalities has been well-studied, the cell circuit mechanisms by which signals from different sensory systems are integrated to control behavior remains poorly understood. Here, we provide a new behavioral model to study the mechanisms of multisensory integration. This behavior, which we termed odor-induced visual valence reversal, occurs when the innate avoidance response to a small moving object by flying *Drosophila melanogaster* is reversed by the presence of an appetitive odor. Instead of steering away from the small object representing an approaching threat, flies begin to steer towards the object in the presence of odor. Odor-induced visual valence reversal occurs rapidly without associative learning and occurs for attractive odors including apple cider vinegar and ethanol, but not for innately aversive benzaldehyde. Optogenetic activation of octopaminergic neurons robustly induces visual valence reversal in the absence of odor, as does optogenetic activation of directional columnar motion detecting neurons that express octopamine receptors. Optogenetic activation of octopamine neurons drives calcium responses in the motion detectors. Taken together, our results implicate a multisensory processing cascade in which appetitive odor activates octopaminergic neuromodulation of visual pathways, which leads to increased visual saliency and the switch from avoidance to approach toward a small visual object.

## INTRODUCTION

Behavioral reactions by animals to environmental sensory stimuli are sometimes reflexive and stereotyped, but can also vary depending on contextual conditions such as locomotor activity state, internal physiological states like hunger or thirst, updated sensory conditions, or signaling from another sensory modality. For example, in walking *Drosophila* behavioral avoidance of an aversive odor can be modulated by food odors (Turner and Ray, 2009) or reversed during flight (van Breugel et al., 2017; Wasserman et al., 2013). Female mosquitoes vigorously approach and explore a small contrasting visual object in the presence of highly attractive carbon dioxide gas, but largely ignore the object in clean air (van Breugel et al., 2017, 2015).

Behavioral plasticity in insects has been broadly attributed to the action of biogenic amines including dopamine, serotonin, histamine and octopamine, which can act as neurotransmitters, neuromodulators or neurohormones (Blenau and Baumann, 2001; Monastirioti, 1999). These derivatives of amino acids exert neuromodulatory effects on behaviors such as olfactory learning, aggression, feeding and egg-laying (Alekseyenko et al., 2014, 2013; Certel et al., 2007; Dierick and Greenspan, 2007; Inagaki et al., 2014, 2012; Keene and Waddell, 2007; Kim et al., 2017; Rezával et al., 2014; Zhou et al., 2012, 2008). As biogenic amines mediate their modulatory effects via second messenger systems, neuromodulation of behavior has generally been shown to occur over minutes to hours (Bargmann, 2012; Marder, 2012). However, those biogenic amines acting like neurotransmitters can exert their functions within seconds (Farooqui, 2012; Monastirioti, 1999). The onset of flight in *Drosophila* modulates the response gain of directionally selective visual motion detecting neurons via octopamine release, which occurs essentially immediately upon the transition from quiescence to flight (Suver et al., 2012). However, the cell circuit mechanisms by which biogenic amines couple different sensory modalities in order to modulate or even reverse behavioral reactions to sensory stimuli remain poorly understood.

Here, we used an electronic visual flight simulator (Reiser and Dickinson, 2008) equipped for odor delivery (Chow and Frye, 2008) to confirm that whereas tethered flies strongly steer toward an elongated bar likely representing a landscape feature, the same flies steer to avoid a small moving square object possibly representing a threat (Maimon et al., 2008). However, we show that flies reverse their behavior to approach when the object is paired with an attractive food odor. We find that this visual valence reversal is odorant-specific: an aversive odor does not reverse object aversion. We show that optogenetic activation of either octopaminergic interneurons or their targets in the visual system is sufficient to elicit visual valence reversal in the absence of odor. Finally, two-photon calcium imaging demonstrates that optogenetic activation of octopaminergic neurons, which are presynaptic in the optic lobe, directly excites columnar motion detecting neurons. Our results suggest a parsimonious model by which odor-activated octopamine release excites visual motion detectors to increase the perceptual salience of a small object, perhaps making a normally aversive small object appear like an attractive feature such as a vertical bar.

## RESULTS

### Visual object valence is reversed by appetitive odor, but not aversive odor

A fly tethered within a wrap-around display of light emitting diodes (LEDs) will steer its wings in response to visual stimuli revolving around the display. Presenting the animal with an elongated vertical bar oscillating in the visual periphery elicits robust wing steering kinematics toward the bar. Reducing the vertical size of the bar to a small box and oscillating this visual object in the same azimuthal location evokes a steering response oriented away from the object (Maimon et al., 2008). For mosquitoes flying freely in a wind tunnel, odor increases the saliency of a small visual object on the floor, causing them to approach and land near the object with greater frequency than in clean air (van Breugel et al., 2015). Thus, we reasoned that olfactory cues might modulate the aversive responses of flies to a small visual object.

We hypothesized that apple cider vinegar (ACV), which is highly attractive to *Drosophila* (Budick and Dickinson, 2006; Duistermars et al., 2009; Semmelhack and Wang, 2009), would attenuate or potentially reverse the behavioral aversion to a small visual object. To test this hypothesis, we utilized an apparatus composed of an LED visual flight simulator equipped with an odor delivery nozzle (Chow and Frye, 2008) and measured fly steering responses to a small object in the presence or absence of ACV (Figure 1A). We adopted an experimental approach similar to that of Maimon et al. 2008, comparing a fly’s steering reactions to a 30-degree square object and a 30×120-degree vertical bar. Steering responses are quantified as the difference in wing beat amplitude (ΔWBA) encoded by an optical wingbeat analyzer. Positive values represent steering torque towards the fly’s right side and negative values steering torque towards its left (Tammero et al., 2004) (Figure 1A’). Prior work (Maimon et al, 2008) used solid black objects set against a uniform white background; these objects are discriminated both by their movement and luminance. To restrict stimulus complexity, we used textured objects set against a textured background to reduce luminance effects (Figure 1B). The two visual objects were oscillated at 1Hz about a point 45-degrees to either side of the visual midline (Figure 1C). To facilitate immediate visual comparison of fly steering direction, we plotted time (in seconds) along the vertical axis and the steering response, ΔWBA, on the horizontal axis such that the plot region to the left of visual midline (green rectangle) corresponds to steering responses towards the left, and the area to the right of midline (gray rectangle) corresponds to turning towards the right.

**Figure 1.**
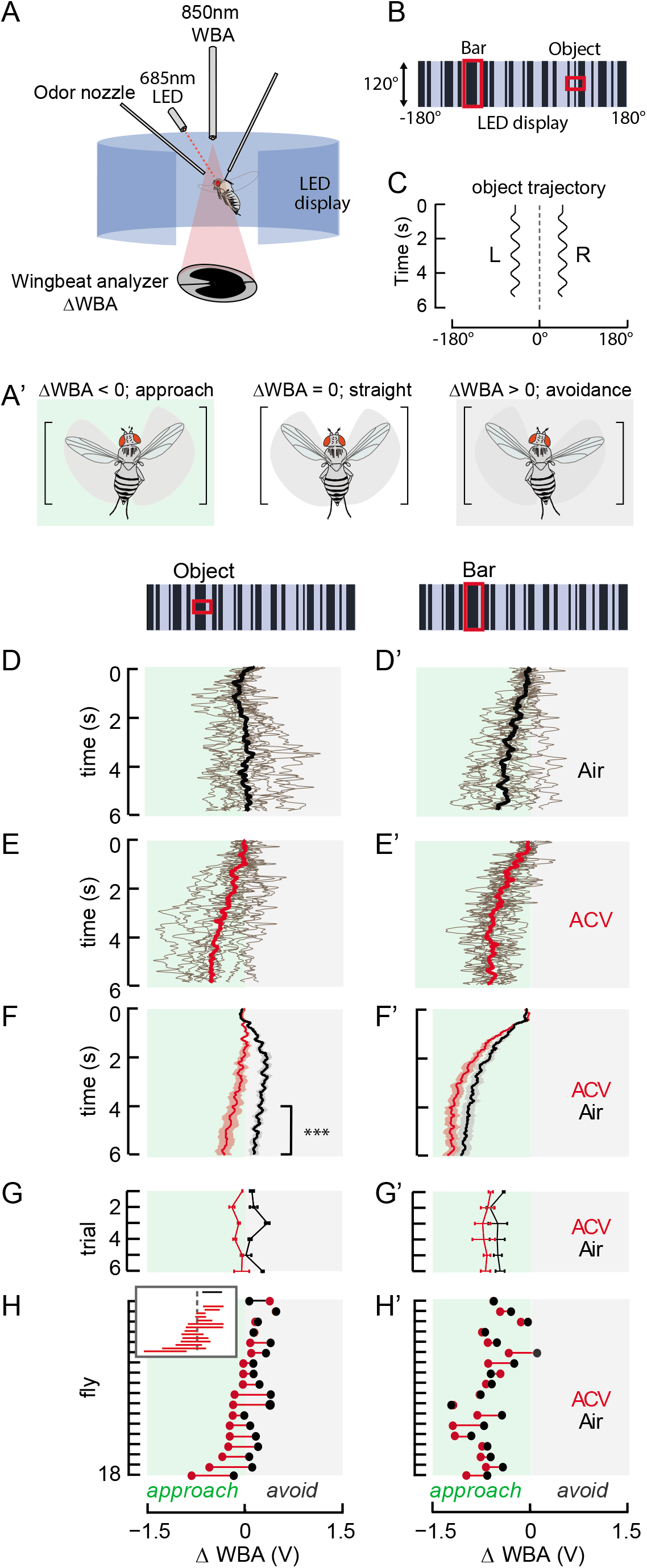
Odor-induced visual valence reversal is odorant specific and learning-independent. (A) Schematic of the rigid tether visual flight arena equipped for either odor delivery or Chrimson optogenetics. The arena is comprised of a series of LED panels controlled in MATLAB. Fly is rigidly tethered onto a tungsten pin and placed between an infrared diode and sensor that measure steering effort. Either an odor port or a LED is placed in front of the fly. For odor experiments, green panels (570nm) were used, and an odor port was placed in front of the fly. For optogenetics experiments, blue panels (470nm) were used to minimize unwanted Chrimson activation. A far-red LED (685nm) was placed in front of the fly. (B) Representation of the visual stimuli used in all flight behavior experiments. A textured, 30×30° square object and 30×120° vertical bar oscillated via a 1Hz sine wave against a textured background. Stimuli moved ±15° in the left or right front quadrants of the arena. (C) Wave form of the 1Hz sine wave used in the odor experiments to oscillate visual stimuli on either the left or right side of the arena. (D&D’) Individual trial ΔWBA responses (gray), calculated as the difference between left and right wingbeat amplitudes (WBA), of a single fly when each visual stimulus was shown in air, superimposed with the mean ΔWBA response across trial (black). By convention, positive ΔWBA denotes steering towards the right and negative ΔWBA denotes steering towards the left. Data are pooled as if all visual stimuli were presented on the left front quadrant of the arena. Therefore, negative ΔWBA (green rectangle) indicates flies are steering to the same side of the visual stimuli, or “approaching” the stimuli, and positive ΔWBA (gray rectangle) represent flies “avoiding” the stimuli. (E&E^1^) Individual trial ΔWBA responses (gray) of a single fly when each visual stimulus was presented with ACV, superimposed with the mean ΔWBA response across trial (red). (F&F^1^): Mean ΔWBA (solid line) and SEM (shaded regions) across wild-type PCF flies (n=18) to an object (D) or bar (D’) in air (black) vs. ACV (red). Bracket denotes the epoch (defined as the last two seconds of the experimental trials) used to calculate mean epoch ΔWBA for statistical analysis and individual fly behavior variability (H). Asterisks denote odor-induced visual valence reversal (p << 0.01, Student’s paired t-test of the last 2 seconds). (G&G’) Mean epoch ΔWBA and SEM of single, consecutive trials, averaged across flies (n=18). (H&H’) Mean epoch ΔWBA of each fly in air (black) vs. ACV (red), sorted by ΔWBA values. Responses from the same fly are joined. The two dots are connected by a red line for steering shifts toward the visual object upon ACV presentation and a black line for ACV shifts away from the visual object. Inset: representation of the same plot, but with dots removed. Red lines indicate that flies tested (12 out of 18) shifted their steering effort toward the visual object when ACV was presented, and black lines indicate that flies (6 out of 18) steered farther away from the visual object in the presence of ACV. Vertical, dashed gray line represents visual midline.

Consistent with Maimon et al. (2008), we demonstrate that in odorless air, the average response of a single wild-type fly indicates subtle steering away from the side of the arena displaying the small object (Figure 1D, thick black line), with varied responses amongst single trials (Figure 1D, thin gray lines). The same animal steers robustly toward a long vertical bar oscillating at the same azimuthal position (Figure 1D’). We observed similar approach or avoidance responses regardless of which side of the arena the visual stimuli were presented (Supp Figure 1). Therefore, for simplicity, trial data in which the stimuli were presented on the right side of the arena were reflected about the midline, pooled with the left side data and plotted as if stimuli were presented on the animal’s left visual field. Remarkably, we found that after switching the odor stream from clean air to ACV, the same animal reverses its behavior and begins to approach the small visual object (Figure 1E), whereas the bar approach response grew stronger in ACV (Figure 1E’). Similar responses were observed for a population of 18 animals, steering to avoid the object and approach the bar in air, but with paired ACV switching from aversion to approach in response to the small object (Figure 1F,F’ p << 0.01, Student’s paired t-test of ΔWBA steady-state mean of the last two seconds).

To examine the variability across animals, we calculated the mean ΔWBA steering responses over the last two seconds of each trial, after responses reached a steady-state, and compared these responses to each visual stimulus before and after ACV presentation for each fly (Figure 1H). Each black dot represents mean steady-state steering response for one animal to the visual trial in air, and each red dot represents the corresponding mean in ACV. The two dots are connected by a red line for ACV reversal to approach in ACV and a black line for shifts further away from the small object. 12 out of 18 flies tested exhibited small object valence reversal (Figure 1H, red lines). The inset shows the same plot, but with the dots removed to provide a compact visual assessment of the phenomenon. For 2 out of 18 flies, the mean steering responses to the small object were not influenced by ACV in either direction (overlap of black and red dots) and were therefore omitted (white space) from the inset. ACV increases the attractiveness of the bar by comparison to the odorless control air stream (p < 0.01, Student’s paired t-test, Figure 1F’). This was consistent across repeated trials (Figure 1G’) and occurred in 10 out of 18 flies tested (Figure 1H’).

This behavioral experiment, with multiple repeated trials for each of two visual stimuli and two olfactory conditions, lasts nearly 10 minutes for each animal. It seemed reasonable that the observed small object tracking behavior could be impacted by visual reinforcement learning, in which a fly might associate the small visual object (“conditioned stimulus”) with a strong, attractive food odorant, ACV (“unconditioned stimulus”), over time (Semmelhack and Wang, 2009). To assess this possibility, we examined how steering responses might change across sequential trials and found that object responses were invariant over the duration of the experiment: flies switch to approach the small object within roughly two seconds after the first presentation of ACV (Figure 1G, trial 1). We found no statistical difference in steering response to the small object between the first or last trial (p = 0.95, trial 1 vs. trial 6, red). Thus, the effect of odor reversing the small object valence persists throughout the experiment, rather than building gradually over time.

We next examined whether visual valence reversal by odor persists across fly strain and odor type. Similar to WT-PCF flies, OregonR flies in clean air steer to avoid the small object and robustly tracks the bar (Figure 2A, black). In the presence of ACV, OregonR flies reverse their steering behavior to approach the small object (p<0.01, Student’s paired t-test of the mean epoch ΔWBA). The reversal was observed in 11 out of 13 flies (Figure 2A, inset). Approach toward the bar was slightly enhanced (p=0.34, Figure 2A’). We next tested a different odorant, ethanol, shown to be attractive in flight (van Breugel et al., 2017). We found that like ACV, EtOH presented to WT-PCF flies acts to reverse the visual valence from object avoidance to approach (Figure 2B, p<0.01). Reversal was observed in 10 out of 13 flies tested (Figure 2B, inset), and the strength of the approach toward the elongated bar was also enhanced (Figure 2B’, p= 0.07). By contrast, when tested with an odorant, benzaldehyde (BA), that flies actively avoid during flight (Wasserman et al., 2012), *Drosophila melanogaster* continue to avoid the small object while continuing to approach the vertical bar in a manner statistically indistinguishable from the responses to odorless air (p=0.20 object; p=0.23 bar, Figure 2C,C’). These results demonstrate that visual valence reversal is odor-context specific, being mediated by olfactory processing circuits conveying at least two commonly appetitive odors, rather than being activated by any odorant.

**Figure 2.**
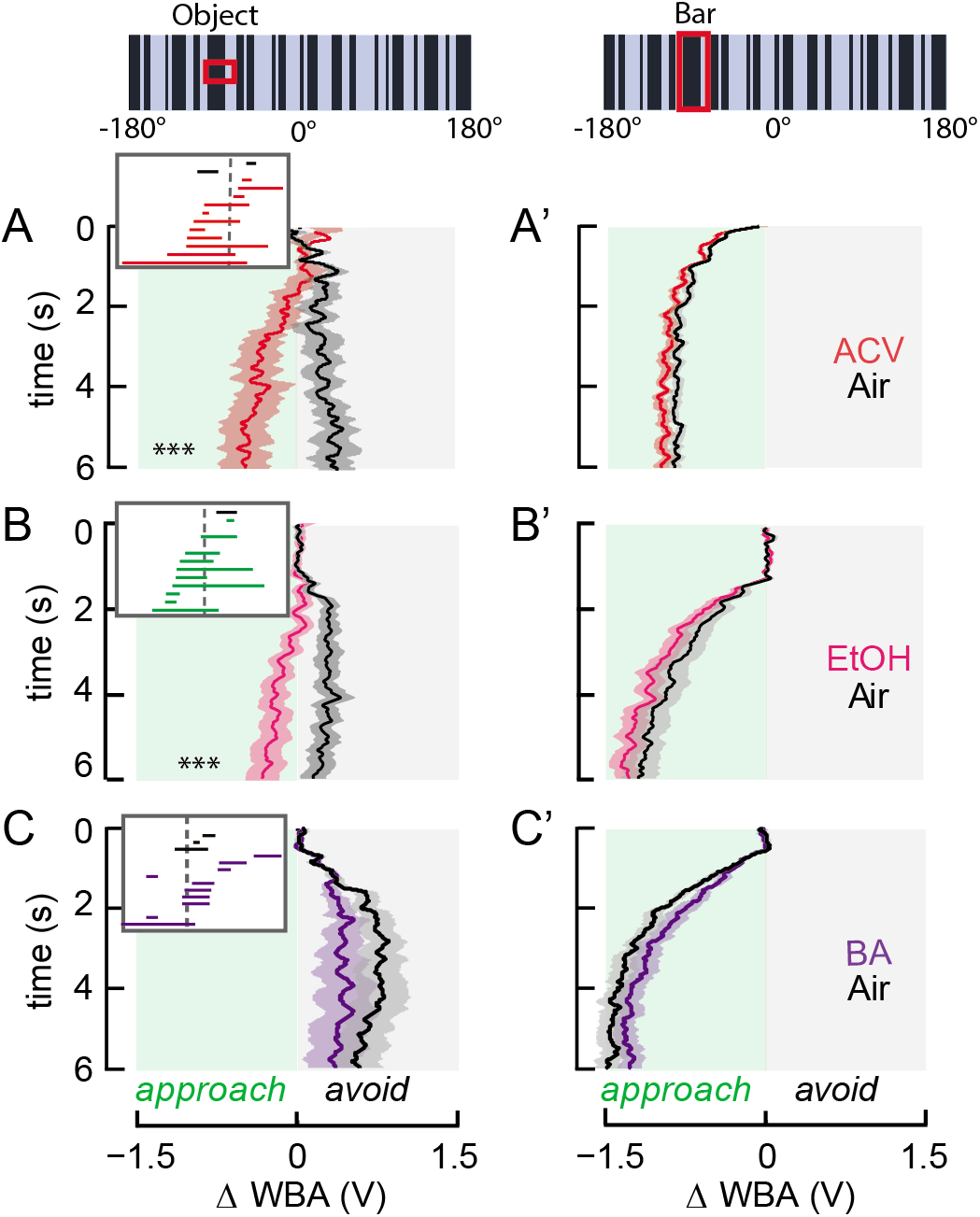
Visual valence reversal is triggered by attractive odors, not aversive odor. (A&A’) Mean ΔWBA and SEM across WT-OregonR flies (n=13) to a small object or bar. Asterisks indicate odor-induced visual valence switch (p<<0.01). Inset: red lines indicate steering shifts toward the visual object upon ACV presentation and black lines represent ACV shifts away from the visual object. Mean steering responses to the small object that were not influenced by ACV in either direction are omitted (empty rows). Vertical, dashed gray line represents visual midline. (B&B’) Mean ΔWBA and SEM across WT-PCF flies (n=13) to an object or bar in the presence of another attractive odorant, ethanol (*** p<<0.01). (C&C’) Mean ΔWBA and SEM across WT flies (n=14) to an object or bar in the presence of an aversive odor, benzaldehyde.

### Optogenetic activation of octopaminergic and tyraminergic neurons induces visual valence reversal

To test whether aminergic neuromodulation is involved in odor-induced visual valence reversal, we expressed Chrimson, a red-shifted excitatory channelrhodopsin (Inagaki et al., 2014; Klapoetke et al., 2014), in candidate aminergic neurons and used a similar experimental paradigm, with the odor delivery system replaced by a 685nm LED (Figure 1A). The inducible nature of Chrimson allows for a within-subject experimental design to directly compare each fly’s flight steering response in the absence (“LED Off”) or presence (“LED On”) of light-activated membrane depolarization. An enhancer-less Gal4 line ‘Empty-Gal4’ (Hampel et al., 2017) served as a genetic control, which evokes behavioral responses to the small object and elongated bar that are similar to those of wild-type flies (Figure 3A, B), although the 685nm LED seems to slightly increase approach toward the bar (Figure 3B’). Optogenetic depolarization of octopaminergic and tyraminergic neurons by the Tdc2-Gal4 driver (Cole et al., 2005) in the absence of odor strongly reverses the steering responses to the small object from aversion to approach, while also increasing the steering responses toward the bar (Figure 3C, Supp Fig 2C, Student’s paired, t-test, *** p << 0.01). 15 out of the 16 flies tested showed valence reversal upon Tdc2>Chrimson activation (Figure 3C, inset, red). This result is observed only with Tdc2-Gal4: optogenetic activation of broadly labeled dopaminergic neurons (p=0.956, TH>Chrimson, Figure 3D), mushroom-body specific dopaminergic neurons (p=0.045, PAM>Chrimson, Figure 3E), or serotonergic neurons (p=0.866, TRH>Chrimson, Figure 3F) failed to evoke visual valence reversal, particularly over the last two second epoch of each trial. The difference in steering amplitude of object responses to LED-ON and LED-OFF by PAM>Chrimson was statistically significant, but activating these neurons merely weakened the small object avoidance and did not reverse it (Figure 3E). Our results indicate that depolarizing the ensemble of tyraminergic and octopaminergic neurons labeled by the Tdc2-Gal4 driver is sufficient to robustly induce visual valence reversal in the absence of appetitive odor (Figure 3C, asterisks).

**Figure 3.**
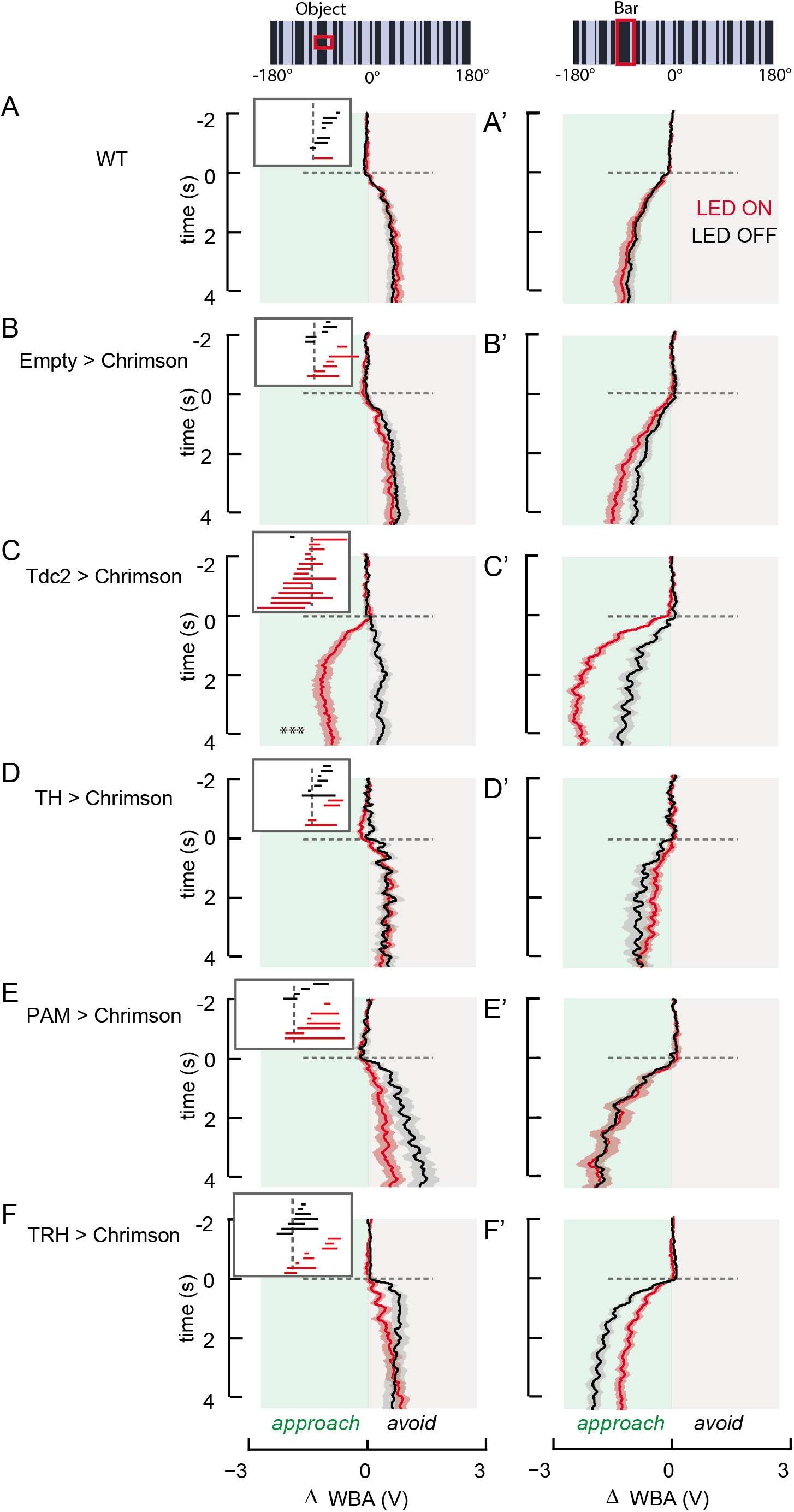
Optogenetic activation of aminergic neurons reveal that OA is sufficient for odor-induced visual valence reversal. Mean ΔWBA (solid line) and SEM (shaded region) to an object (A-F) or bar (A’-F’) in LED Off (black) or LED On (red) conditions. Each panel row represents flies of one distinct genotype. Horizontal dashed line represents the onset of visual stimulus motion. Green rectangles represent portion of ΔWBA values that correspond to when flies are approaching the visual stimulus. Asterisks (C) indicates visual valence switch (p << 0.01, Student’s paired t-test of the last 2 seconds). Inset denotes whether each fly (each line along the vertical axis), sorted by ΔWBA values, steer more towards (red) or away from (black) the object in the presence of ACV. Mean steering responses to the small object that were not influenced by ACV in either direction are omitted (empty rows) (A, A’) n=11; (B, B’) n=12; (C, C’) n=16; (D, D’) n=13; (E, E’) n=12; (F, F’) n=16.

### Optogenetic activation of T4/T5 motion detectors induces visual valence reversal

To begin to address the underlying mechanism of action by Tdc2 neurons, we assessed their synaptic organization in the optic lobe by co-labeling with DenMark (Nicolaï et al., 2010) and synaptotagmin (Zhang et al., 2002). Consistent with previous findings (Busch et al., 2009), we found that Tdc2 neurons are broadly presynaptic in the optic lobe, showing strong and broadly distributed *syt* labeling throughout the medulla and lobula complex (Figure 4A). We also noted thicker and less dense axonal (DenMark) projections returning to the central brain and heavy labeling in the sub-esophageal zone.

**Figure 4.**
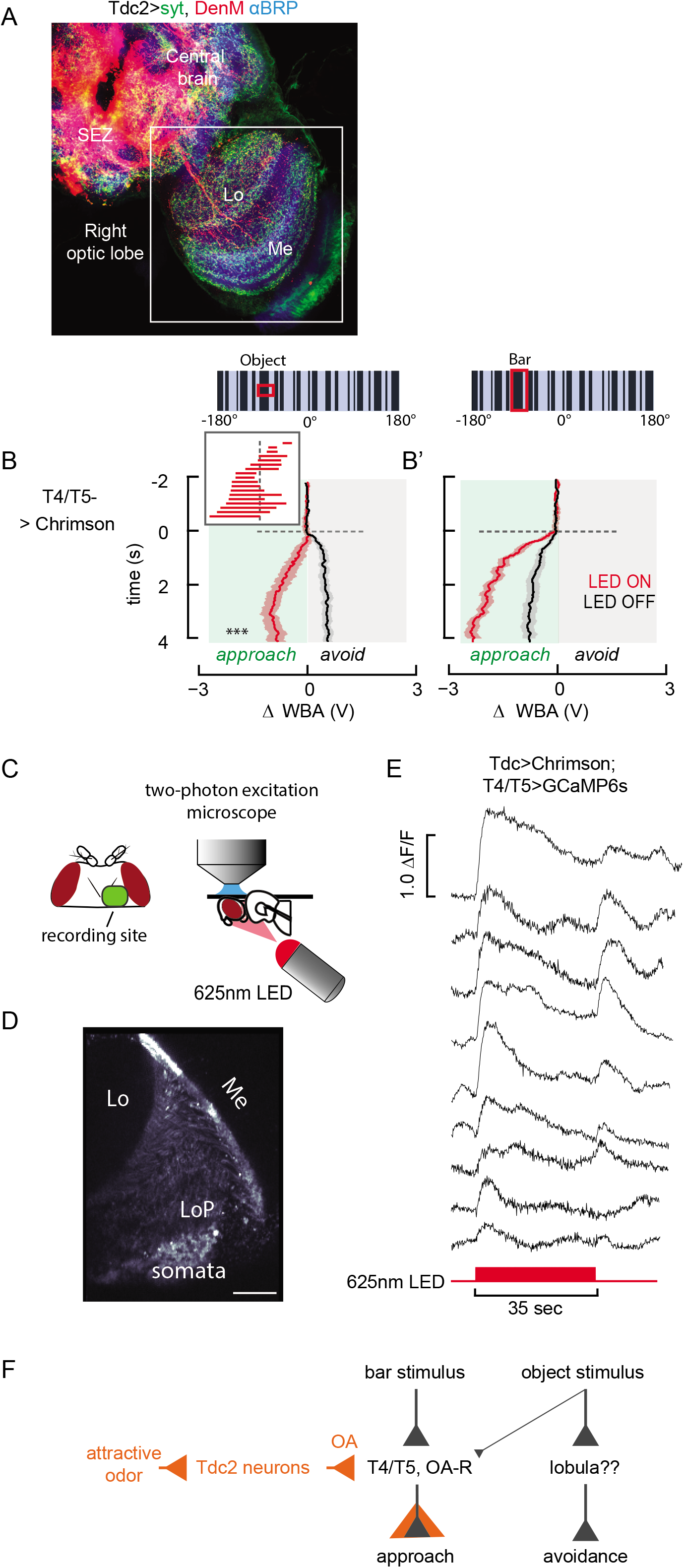
Optogenetic activation of T4/T5 motion detectors triggers visual valence reversal. (A) Distribution of presynaptic (green) and postsynaptic (red) neurites of the Tdc2-Gal4 driver. Red = DenMark labeling; Green = synaptotagmin labeling; Blue = anti-BRP labeling. (B & B’) Mean ΔWBA (solid line) and SEM (shaded region) to an object (B) or bar (B’) in LED Off (black) or LED On (red) conditions. Optogenetic activation of T4/T5 neurons (n=18) is sufficient to phenocopy odor-induced visual valence reversal (C, asterisks, p << 0.01, Student’s paired t-test of the last 2 seconds). An enhanced bar tracking behavior was also observed upon Chrimson excitation (B’, p << 0.01). (C) Schematic of the two-photon *in vivo* calcium imaging set-up. A quiescent fly is fixed and the back of the head is opened to allow optical access. A 625nm red LED is placed behind the fly and illuminates the entire fly during the LED On condition. (D) Example of an imaging plane and the visible T4/T5 region. For our analysis, a polygon is drawn (not shown) around the visible T4/T5 neurons and analyzed in its entirety. (E) ΔF/F of the background subtracted T4/T5 in vivo calcium response before, during and after Tdc2 activation with the red LED. Each trace is data from a single fly preparation (n=9). (F) Summary diagram of olfactory neuromodulation of visual valence reversal. Lines with filled triangle arrows represent excitatory interactions. Thickness of the lines and triangles indicates interaction strength. Orange indicates odor ON. Two parallel visual channels control bar approach and object avoidance. Object stimuli weakly drive T4/T5, where response gain is increased by odor. With odor (or optogenetic activation of upstream components) object stimuli drive approach responses more powerfully than aversion responses.

We next sought to reveal the visual circuits involved in visual valence reversal behavior. Columnar T4 and T5 neurons of the medulla show directionally selectivity, express OA receptors (Davis et al., 2018; Pankova and Borst, 2016), and are presynaptic to wide-field neurons of the third optic ganglion that exhibit Tdc2-dependent increase in visual response gain upon flight initiation (Suver et al., 2012), one of which has been shown to be modulated by odor (Wasserman et al., 2015). We thus reasoned that Tdc2>Chrimson rapidly releases OA to enhance synaptic drive within motion T4/T5 detectors. To test this hypothesis, we expressed Chrimson in T4/T5 neurons and subjected these flies to a similar optogenetics behavioral paradigm, but without the 2-minute priming excitation used in the Tdc2>Chrimson experiment. We found that optogenetically activating T4/T5 neurons with a 685nm LED at 0.010 mW/mm^2^ intensity mimicked the influence of both appetitive odor and Tdc2>Chrimson, increasing the strength of steering responses toward the bar and reversing the steering responses from aversion to approach toward to the small object (p << 0.01, Figure 4B, B’). This was observed in every of the 18 flies tested (Figure 4B, inset). Since T4/T5 neurites innervate four layers of the lobula plate that each represent a separate cardinal direction of motion (Maisak et al., 2013; Mauss et al., 2014), one might expect that optogenetically depolarizing the full population of directionally tuned T4 and T5 small-field motion detecting neurons would render flies unable to perceive and respond to directional motion cues. Indeed, when we increased the LED intensity 4-fold to 0.040 mW/mm^2^, we observed weakened approach behavior to both the small object and vertical bar (Supp Figure 3B&B’, yellow). We also observed weaker wide-field optomotor responses at the higher LED intensity compared to the lower LED intensity (Supp Figure 3B”). These results suggest that mild T4/T5 depolarization is sufficient to induce visual valence reversal from avoidance to approach toward the small object (Figure 4B), while strengthening approach toward the bar (Figure 4B’), responses that are qualitatively similar to the effects of ACV or EtOH (Figure 1F,F’, 2B,B’). These results also demonstrate that appropriately tuned Chrimson excitation can enhance the synaptic drive from motion detection circuitry.

### T4/T5 motion detectors are excited by Tdc2 neurons

Activating octopaminergic neurons that are presynaptic in the optic lobe via Tdc2>Chrimson phenocopied appetitive odor delivery to reverse the behavioral valence of a small object in flight (Figure 3C). So did activating T4/T5 motion detectors whose inputs are broadly excited by OA and themselves express OA receptors (Figure 4B)(Pankova and Borst, 2016). Prior results showed that Tdc2 activity is necessary and sufficient for locomotion-induced gain increase in lobula plate tangential cells (Suver et al., 2012), and appetitive odor (ACV) can elicit Tdc2 calcium responses in a quiescent preparation (Wasserman et al., 2015). Thus, we sought to test whether Tdc2 neurons functionally interact with T4/T5 motion detectors.

We expressed Chrimson in Tdc2 neurons and GCaMP6s in T4/T5 neurons. Using two-photon excitation microscopy, we imaged calcium accumulation in T4/T5 processes before and after Chrimson excitation *via* an external 625nm LED (Figure 4C). Each experiment was performed only once on each fly and comprised of recording baseline T4/T5 activity followed by simultaneous Tdc2>Chrimson activation and T4/T5>GCaMP6s imaging. For each recording, we selected any active T4/T5 neurites visible in our imaging plane (Figure 4D) as our ROI for analysis. We observed increases in T4/T5 calcium activity immediately upon Chrimson excitation of Tdc2 (Figure 4E). These calcium responses were characterized by an onset transient, decaying steady-state response, and a secondary calcium transient after the LED is switched off (Figure 4E). Qualitatively similar rapid onset T4/T5 excitatory responses to optogenetic depolarization of Tdc2 neurons were observed for each fly tested.

## DISCUSSION

In this study we characterized a novel behavior of *Drosophila melanogaster*, odor-induced visual valence reversal. During both free and tethered flight, *Drosophila melanogaster* tend to approach an elongated vertical edge likely representing a landscape feature such as a plant stalk (Fox et al., 2014; Frye et al., 2003; Maimon et al., 2008; van Breugel and Dickinson, 2012), while at the same time steering to avoid a small moving object, presumably representing a threat such as an approaching predator (Maimon et al., 2008). Small object aversion has been demonstrated in freely walking and freely flying *Drosophila melanogaster* (Grabowska et al., 2018; Maimon et al., 2008). However, closed-loop object avoidance by rigidly tethered flies is characterized by a weakened frontal fixation response, rather than robust anti-fixation (Maimon et al., 2008). We have obtained similar results (data not shown). Innate object avoidance behavior by a rigidly tethered fly is most robust and amenable to experimental manipulations under open-loop feedback conditions, which we have focused on here to maximize our measurement fidelity.

We showed that the avoidance response to a small moving object is not a rigid fixed action pattern, but is instead modulated by sensory state, switching from avoidance to approach in the presence of appetitive food odors such as apple cider vinegar - ACV (Figure 1F) or ethanol – EtOH (Figure 2B), but not of an aversive odor, benzaldehyde - BA (Figure 2C). This odor-induced visual valence reversal does not appear to be learning-dependent (Figure 1G). Optogenetic activation of Tdc2-labeled octopaminergic neurons, but not dopaminergic nor serotonergic neurons, phenocopies the effect of ACV and EtOH on visual valence reversal (Figure 3C). OA has been shown to increase the strength of visual responses throughout the cellular motion vision pathway of the optic lobe (Maimon et al., 2010; Suver et al., 2012; van Breugel et al., 2014), where Tdc2 neurons have both pre- and post-synaptic processes (Figure 4A)(Busch et al., 2009). Remarkably, optogenetic activation of small-field motion detecting T4/T5 neurons also robustly induces small object valence reversal (Figure 4B). Finally, optogenetic activation of octopamine increases GCaMP fluorescence in T4/T5 in the absence of any visual stimulus (Figure 4E). Taken together, our results provide a novel model for multisensory processing in which an appetitive odor stimulates octopamine release that increases the response gain of the motion vision pathway to influence object tracking behavior. Our results do not preclude the participation of other visual pathways, especially those that are involved in small object aversion. It is largely unknown how the motion vision and object vision pathways interact. We also don’t yet understand how OA neurons of the optic lobe are stimulated by the olfactory system.

### Odor-activated visual valence reversal is rapid and odor-valence specific

Although we have not tested an exhaustive panel of odorants, our work to date suggests that visual valence reversal is mediated by appetitive but not aversive odors, a result that provides key insight into the putative organization of olfactory inputs to OA neuromodulation. In the *Drosophila* olfactory system, diverse lines of evidence suggest that attractive and aversive odorants are encoded by anatomically segregated pathways that reside within subdomains of hierarchical olfactory neuropils (Grabe and Sachse, 2018; Masse et al., 2009; Sachse and Beshel, 2016; Schultzhaus et al., 2017). Our finding that the observed odor-induced visual valence reversal takes place only in the presence of attractive odorants (ACV and EtOH, Figure 1F & 2B) and not aversive odorants (BA, Figure 2C) specifically implicates the attractive olfactory pathway (Masse et al., 2009; Sachse and Beshel, 2016; Strutz et al., 2014).

A third-order olfactory neuropil, the mushroom body (MB), is classically known for its role in olfactory learning (de Belle and Heisenberg, 1994) in flies, yet is recently receiving attention for its role in behavioral plasticity (Grunwald Kadow, 2018). For example, the MB is required to encode the innate valence of odors, independent of learning (Bräcker et al., 2013; Lewis et al., 2015). The MB has been shown to play a role in various visual behaviors (Aso et al., 2014; van Swinderen, 2009; Vogt et al., 2016, 2014). Rapid odor modulated optomotor reflexes require intact MB circuitry (Chow et al., 2011). We have found no compelling evidence to implicate the MB in visual object valence reversal, which is learning-independent in that it has a rapid onset and does not improve with time (Figure 1G). Additionally, optogenetic activation of PAM neurons, a subset of DA neurons that modulate the synapse between MB Kenyon Cells (KCs) and output neurons (MBONs) in olfactory reward learning (Hattori et al., 2017; Hige et al., 2015; Hige and Turner, 2015; Keene and Waddell, 2007; Liu et al., 2012; Vogt et al., 2014) was not sufficient to phenocopy odor-induced valence reversal (Figure 3E).

Another third-order olfactory neuropil, the lateral horn (LH), mediates olfactory behaviors in a rapid, experience-independent manner (Fişek and Wilson, 2014; Grabe and Sachse, 2018; Sachse and Beshel, 2016; Schultzhaus et al., 2017). This neuropil houses neurons that encode odor features such as hedonic valence and odor intensity (Grabe and Sachse, 2018; Sachse and Beshel, 2016; Schultzhaus et al., 2017). LH projection neurons innervate the superior medial protocerebrum (SMP), superior lateral protocerebrum (SLP), and the ventral nerve cord (VNC) (Dolan et al., 2018; Fişek and Wilson, 2014; Jefferis et al., 2007; Sachse et al., 2007; Tanaka et al., 2012, 2004). To our knowledge, there are as yet no known projections between MB or LH neuropils to Tdc-2 labeled neurons. However, LHNs receive input from the sub-esophageal zone, where Tdc2 is heavily dendritic (Figure 4A)(Busch et al., 2009; Cole et al., 2005; Dolan et al., 2018). Although ACV has been shown to excite Tdc2 neurons in the optic lobe (Wasserman et al. 2015), the olfactory inputs to Tdc2 neuromodulatory circuits of the brain remain unknown.

### Odor-induced visual valence reversal is likely mediated by octopamine

The Tdc2-Gal4 driver used in our experiments labels both octopaminergic and tyraminergic (TA) neurons (Cole et al., 2005). Work in the fly field frequently discusses Tdc2-Gal4 and OA neurons synonymously, yet we must be careful not to rule out a potential role of TA neurons in mediating odor-induced visual valence reversal. However, several lines of evidence suggest that OA, not TA, is mediating the effects we see. OA and TA generally act through independent G-protein coupled receptors (with the exception of an octopamine-tyramine receptor), and exert opposite effects on cAMP and Ca^2+^ levels downstream of the GPCR (Roeder, 2005). Thus, OA and TA often have antagonistic effects (Saraswati et al., 2004). We attempted to separate the effects of OA from TA by using TBH^nM18^ mutants, which lack OA and have an elevated level of TA (Monastirioti et al., 1996). However, in our hands these flies exhibited severe flight impairments, consistent with previous reports (Brembs et al., 2007). Another strong implication for OA rather than TA is that all stages of motion vision processing tested to date have been shown to be gain-modulated by chlordimeform, which is an OA-specific agonist (Arenz et al., 2017). Finally, T4/T5 neurons express OA-R’s and few TA-R’s (Pankova and Borst, 2016). Given the available evidence, the most parsimonious model is that odor-induced visual valence reversal is mediated by OA rather than TA.

Octopamine mediates neuromodulation of many behaviors, including aggression, courtship, and egg-laying (Farooqui, 2012, 2007). Our data suggest that OA is also sufficient in mediating odor-induced visual valence reversal. We attempted several experiments to determine the effect of OA inactivation on this behavior. However, these loss-of-function genotypes either did not have a reliable genetic control that allowed us to draw any interpretation or exhibited severe flight defects that rendered the flies unsuitable for our behavioral paradigm. This difficulty is not unexpected, however, as Tdc2-labeled cells represent a widely diverse population of roughly 100 interneurons that innervate the brain and ventral nerve cord (Busch et al., 2009; Sinakevitch and Strausfeld, 2006).

Despite its numerous functional roles, the mechanism by which OA is activated by upstream systems or behavioral states and how it in turn modulates the visual system remain largely unknown. One open question is how the olfactory system is connected with octopaminergic Tdc2-Gal4 labeled cells. Our lab previously provided evidence that Tdc2 calcium responses in quiescent adult flies increased upon the presentation of ACV (Wasserman et al., 2015). Other work has demonstrated that Tdc2 neurons in larvae are activated upon optogenetic excitation of Orco (Ma et al., 2016), a broadly expressed olfactory co-receptor (Larsson et al., 2004). Despite these efforts, precisely which brain regions and neuronal subdomains of Tdc2 neurons in adult flies are activated by attractive odorants, and in turn stimulate the motion vision pathway, requires further investigation.

### Octopaminergic neuromodulation of visual processing is hierarchical

The transition from quiescence to flight or walking behavior in *Drosophila* is associated with increased response gain of and shifted frequency tuning by directionally selective motion detecting neurons of the lobula plate (Arenz et al., 2017; Chiappe et al., 2010; Jung et al., 2011; Maimon et al., 2010; Suver et al., 2012). This modulatory influence by locomotor state has been shown to depend upon the activity of octopaminergic Tdc2-Gal4 neurons (Maimon et al., 2010; Suver et al., 2012). We posit that odor-evoked octopaminergic modulation of visual object valence behavior implicates a hierarchy of OA neuromodulation, because this behavior is triggered by sensory cues encountered during active flight, when the OA neuromodulation on motion vision driven by the onset of locomotion has already occurred. In other words, OA modulation of motion vision circuitry triggered by the onset of active flight is insufficient to trigger visual valence reversal (Figure 1, black). It is the additional presentation of either an appetitive odorant or Tdc2 optogenetic activation in a flying animal that induces visual valence reversal of a small object (Figure 1F & 2C, red). Such a hierarchy could explain why ACV failed to modulate visual responses by T4/T5 neurons in a quiescent imaging preparation (Wasserman et al., 2015).

Hierarchical OA neuromodulation could be mediated by the recruitment of different subpopulations of Tdc2 neurons, or by differences in the distribution and molecular action of the various OA receptor types, or both (Farooqui, 2012). Given the diverse neuronal morphologies within the ensemble of Tdc2-Gal4 neurons (Busch et al., 2009; Claßen and Scholz, 2018), it seems highly unlikely that all OA neurons function as a single ‘mega-interneuron’. Indeed, recent work has shown that, depending on experimental parameters, activation of Tdc2 neurons can decrease song behaviors (O’Sullivan et al., 2018) or promote male-to-male courtship (Certel et al., 2007), demonstrating the functional diversity of Tdc2 neurons and OA action. The extent to which hierarchical OA activation is mediated by either spatial effects via the recruitment of different Tdc2-Gal4 cell types and distribution of different OA receptor types, or by superposed effects by behavioral state and odor within individual neurons, requires further investigation.

### T4/T5 small-field motion detectors support bar tracking behavior

Some sort of Tdc2-dependent octopaminergic modulation of T4/T5 neurons would be expected since T4/T5 neurons express OA receptors (Davis et al., 2018; Pankova and Borst, 2016) and show shifted frequency tuning in response to bath applied chlordimeform (Arenz et al., 2017). Columnar neurons of the medulla presynaptic to T4/T5 neurons also show elevated visual response gain in bath application of chlordimeform (Arenz et al., 2017) or optogenetic activation of Tdc2 (Strother et al., 2018). Our findings that Tdc2 optogenetic activation increases calcium responses of T4/T5 neurons is, to our knowledge, the first evidence for rapid functional interactions between aminergic neuromodulatory cells and these directionally-selective visual processing neurons (Figure 4), as previous findings focused on cells presynaptic to T4/T5.

Chrimson activation of Tdc2 neurons results in peak GCaMP6s fluorescence by T4/T5 cells within 1.5 seconds (Figure 4E), which is in good agreement with the roughly 2-second onset for visual valence reversal behavior (Figure 1F). Our working model is that olfactory or optogenetic excitation of Tdc2 neurons releases OA that binds OA-receptors on T4/T5 neurons, depolarizing them and thereby enhancing the impact of presynaptic drive. Synaptic facilitation would result in larger amplitude responses to either a small moving object or bar by downstream motion integrators during active flight. In support of this model, whereas low-intensity optogenetic excitation of T4/T5 is sufficient to reverse the response to a small object from avoidance to approach (Figure 4B, Supp Figure 3B), stronger excitation of T4/T5 diminishes the magnitude of optomotor responses as might be expected from saturating the responses of all directional motion detectors (Supp Figure 3B).

How does the T4/T5 system support bar tracking behavior in flies? Flies walking on an air supported ball with genetically silenced T4/T5 neurons show strongly attenuated wide-field optomotor responses, but turns to approach a vertical bar were largely intact, suggesting that at least some aspects of bar detection occurs via neural pathways other than small-field T4/T5 motion detectors (Bahl et al., 2013). By contrast, flies in tethered flight under high-gain closed-loop feedback conditions show strongly attenuated ability to frontally fixate a bar when T4/T5 synaptic output is blocked (Fenk et al., 2014). Taken together, these results suggest that directional motion detection by T4/T5 cells contributes some but not all aspects of bar tracking behavior.

By contrast to directional motion vision, the neural pathways for position and non-directional small object motion are poorly understood. In house flies, a class of neurons of the lobula plate are excited by motion within a small receptive field and are inhibited by wide-field displacement of the panorama (Egelhaaf, 1985; Liang et al., 2008) but as yet these cells are unknown in *Drosophila*. Recent studies of the *Drosophila* lobula have characterized several classes of columnar projection neurons that encode visual features such as looming (Namiki et al., 2018), movement of small contrasting targets (Keleş and Frye, 2017), and optical disparities generated by vertical edges moving against a visual panorama that correlate with behavioral reactions to similar single-edge stimuli (Aptekar et al., 2015). How these or other cell classes outside of the T4/T5 directional motion detection pathway participate in object behavior remains to be fully understood.

We propose that the weak synaptic drive that a small visual object would produce in small-field T4/T5 cells essentially goes unnoticed by downstream integration pathways that supply the motion dependent component of bar approach behavior (Figure 4F). Parallel circuitry, possibly through the lobula, would mediate avoidance behavior. Olfactory neuromodulation by octopamine effectively increases the saliency of the small object within the T4/T5 motion pathway, thereby driving a robust bar-like approach response that overrides the subtle aversive response mediated outside of the T4/T5 system (Figure 4F). That the bar responses are boosted by odor supports this conceptual model. Enhanced T4/T5 activity induced by OA alters the synaptic weights between the neural pathways that elicit aversion and approach, effectively reversing avoidance to approach behavior without the requirement of any sort of neural switch (Figure 4F). This simple algorithm allows for modulating the decision processes required to navigate landscape features and also orient toward potential food resources whilst maintaining adaptive avoidance responses to potential threats.

## MATERIALS AND METHODS

### Fly stocks / Fly maintenance

All flies were maintained in a humidity-controlled environment on a 12:12 hour circadian light:dark cycle. Crosses involving aminergic cell types were raised at 18°C (Tritech Research) to reduce Gal4 toxicity. All other flies were raised in a 25°C animal room. For behavior experiments, female flies 3-5 days post-eclosion are used, and experiments are conducted within 4 hours from light on or 4 hours before light off.

The following fly stocks were used: wild-type PCF flies, wild-type OregonR flies, Tdc2-Gal4 (BDSC#9313), TH-Gal4 (BDSC #8848), TRH-Gal4 (gift from Birman lab), Empty-Gal4 (pBDPGal4U, Janelia), T4/T5 split-Gal4 (Janelia SS00324), R42F06-lexA (BDSC#54203), 20xUAS-GCaMP6m (BDSC#42748, 42750), 20xUAS-GCaMP6s(BDSC#42749), 20xUAS-Chrimson::tdTomato-VK5 (gift from Anderson lab, Caltech), w^1118^; UAS-DenMark, UAS-syt.eGFP; D^1^/TM6C (BDSC #33064), 13xLexAop-opGCaMP6s,10xUAS-Chrimson88 (gift from Reiser lab).

### Rigid tether flight simulator and odor delivery

The rigid tether visual arena is previously described (Maimon et al., 2008). The arena comprises of computer-controlled 96×32 cylindrical array of 570nm LEDs, each pixel subtending 3.75 degrees on the retina. Experimental flies were cold-anesthetized and tethered to 0.1mm-diameter tungsten pins and were allowed to recover for one hour after tethering within a plastic box containing a dish of water and illuminated by a heat lamp to maintain humidity. In the flight arena, an infrared emitter and sensor are placed above and below the tethered fly to capture a shadow of the beating wings on the sensor. The sensor and associated electronics measure the amplitude of each wing beat. The difference in amplitude of the left and right wing signals (ΔWBA) is proportional to yaw torque (Tammero, Frye, and Dickinson 2004) and indicates the fly’s attempt to steer left (ΔWBA<0) or right (ΔWBA>0).

The following visual stimuli are used in all behavioral paradigms: a 30 degree, randomly textured square, and an elongated, textured bar 30 degrees in width and 120 degrees in height. Visual stimuli are presented in random order on the left or right 45-degrees from midline. The stimuli oscillate via 1Hz sine wave with an amplitude of 15 degrees. Each experimental condition (object/bar, arena left/right) are presented 4-6 times. Trials in clean air are presented before the odor-paired trials rather than being interspersed in order to limit potential effects of olfactory working memory (Saxena et al., 2017). A closed loop bar fixation trial is placed between open-loop test trials to keep the fly actively engaged in the assay.

Odors used in this study were apple cider vinegar (Ralph’s Grocery generic brand), ethanol (Decon Laboratories, Inc.) diluted to 70% and benzaldehyde (Sigma Aldrich, B1334) diluted to 40%. Because benzaldehyde precipitates easily, the odorant is placed on filter paper inside the odor delivery tube (Wasserman et al., 2012). Odor delivery to the tethered fly has been described previously (Chow and Frye, 2008). Briefly, saturated odor vapor was delivered through a pipet tip placed 1 cm in front of the fly’s head and drawn away by vacuum in a tube positioned behind the fly. To confirm that each fly responded to the odor, we administered a 5-second odor response test without visual cues, and only included flies in the experiment that showed a significant increase in wingbeat frequency upon the onset of the odor pulse (Chow, Theobald, and Frye 2011). No flies were run more than once. After each experiment, a photoionization detector (miniPID Model 200B, Aurora Specific), was used to confirm the ON/OFF switching of air/odor at the location of the tethered fly.

### Rigid flight simulator and optogenetic activation

Optogenetics flight experiments are conducted in a similar setup to the odor experiments, except that blue LED panels (470nm) are used instead of green LED panels (570nm) to avoid Chrimson activation by the display (Klapoetke et al. 2014). The odor delivery system is replaced by a red LED (685nm) that illuminates the entire fly. Similar to the odor experiment paradigm, all LED-Off trials were conducted first, followed by the block of LED-On trials. The same visual patterns from the odor experiments were used and presented with a random block experimental design.

For flies expressing aminergic drivers, a 2-minute closed loop fixation period with the LED On is placed between the LED Off and LED On blocks. This was done to account for the slow kinetics of activation of biogenic amines reported in the literature, which predominantly mediate their effects via G-protein coupled receptors (Monastirioti et al. 1995). For flies expressing Chrimson under the control of T4/T5-splitGal4, the 2-minute ‘preincubation’ LED On period is removed, and the LED is turned off in between trials during the LED On block. Except for the high-intensity experiment (Supp Figure 3B-B”), all optogenetics experiments used a LED power intensity of 0.010 mW/mm^2^. Power intensity was increased to 0.040 mW/mm^2^ for the high-intensity experiment.

### All-trans-retinal

For proper Chrimson protein conformation, all-trans-retinal (ATR, Sigma Aldrich, R2500) is required. Though flies endogenously produce retinal, additional ATR is added to the food to boost performance (Simpson and Looger, 2018). F1 Chrimson flies are raised in 0.5mM ATR food post-eclosion for at least 3 and no more than 5 days before being used for experiments.

### Calcium Imaging

*In vivo* calcium imaging experiments were performed in the setup described in Figure 4D. A red LED (625nm, Thor Labs M625L3) is placed behind the fly and illuminates the entire animal. Baseline responses were recorded first before the LED is switched on. Recordings are taken from the entire T4/T5 population including medulla dendrites, axons, and terminals in the lobula and lobula plate, and GCaMP6s fluorescence is measured on each fly for both baseline LED-Off and LED-On conditions. Preparation for calcium imaging was previously described in (Keleş and Frye, 2017). Briefly, flies were anesthetized on a cold block, and then placed in a hole cut from a stainless steel shim. The head-capsule and thorax were secured to the edges of the hole using UV-activated glue. The proboscis, antennae and legs were immobilized using beeswax. We used a sharp forcep (Dumont, #5) to cut open the left side of the fly head, and removed muscle and fat to enable access to the optic lobe. The prep was constantly bathed in saline solution (103mM NaCl, 3mM KCl, 1.5mM CaCl2, 4mM MgCl2, 26mM NaHCO3, 1mM NaH2PO4, 10mM trehalose, 10mM glucose, 5mM TES, 2mM sucrose).

### Image Analysis

After exporting each recording as a TIFF stack, the images were first aligned to account for any movement shift during recording. T4/T5 ROIs were based on all the visible neurites labeled by GCaMP6s. Background ROI was selected as any patch on the imaging plane that fell outside of the T4/T5 ROI. For each preparation, we recorded 7s of baseline T4/T5 activity, followed by 35s of simultaneous Tdc2 optogenetic activation and T4/T5 GCaMP6s imaging. The raw responses of the background ROI was subtracted from the T4/T5 response, before normalizing to F_0_,defined as the mean response prior to LED on and subtracting 1.

### Statistics

Student’s paired t-test of the last two seconds was performed in MATLAB 2017a (MathWorks, Inc.) to compare mean epoch ΔWBA.

## ACKNOWLEDGMENTS

This work was supported by a grant from the National Science Foundation IOS-1455869. We thank Dr. Ben Hardcastle for help with imaging data analysis, Dr. Mehmet Keleş for help with generating fly lines, and Dr. Sufia Sadaf for help with the confocal image.

## AUTHOR CONTRIBUTIONS

Conceptualization, M.A.F. and K.Y.C.; Methodology, M.A.F. and K.Y.C.; Investigation, K.Y.C.; Writing-Original Draft, K.Y.C.; Writing-Review, Revision & Editing, M.A.F.; Funding Acquisition, M.A.F.; Resources and Supervision, M.A.F.

## SUPPLEMENTAL TEXT

We presented the visual stimuli on both the left and right side of the arena. In the main text, we pooled our data as if visual stimuli were all presented on the left side, since we observed similar behavioral reactions (bar tracking, small object avoidance in air, and small object approach in appetitive odor or LED On) regardless of which side the stimuli was presented. The caveat of pooling the data, however, is that the sinusoidal component of the WBA responses plotted in the supplementary figures were masked since most of our responses were slightly out of phase with each other. The DC component of the responses were preserved, however, and allowed us to make our arguments while presenting our data with visual simplicity.

**S1.**
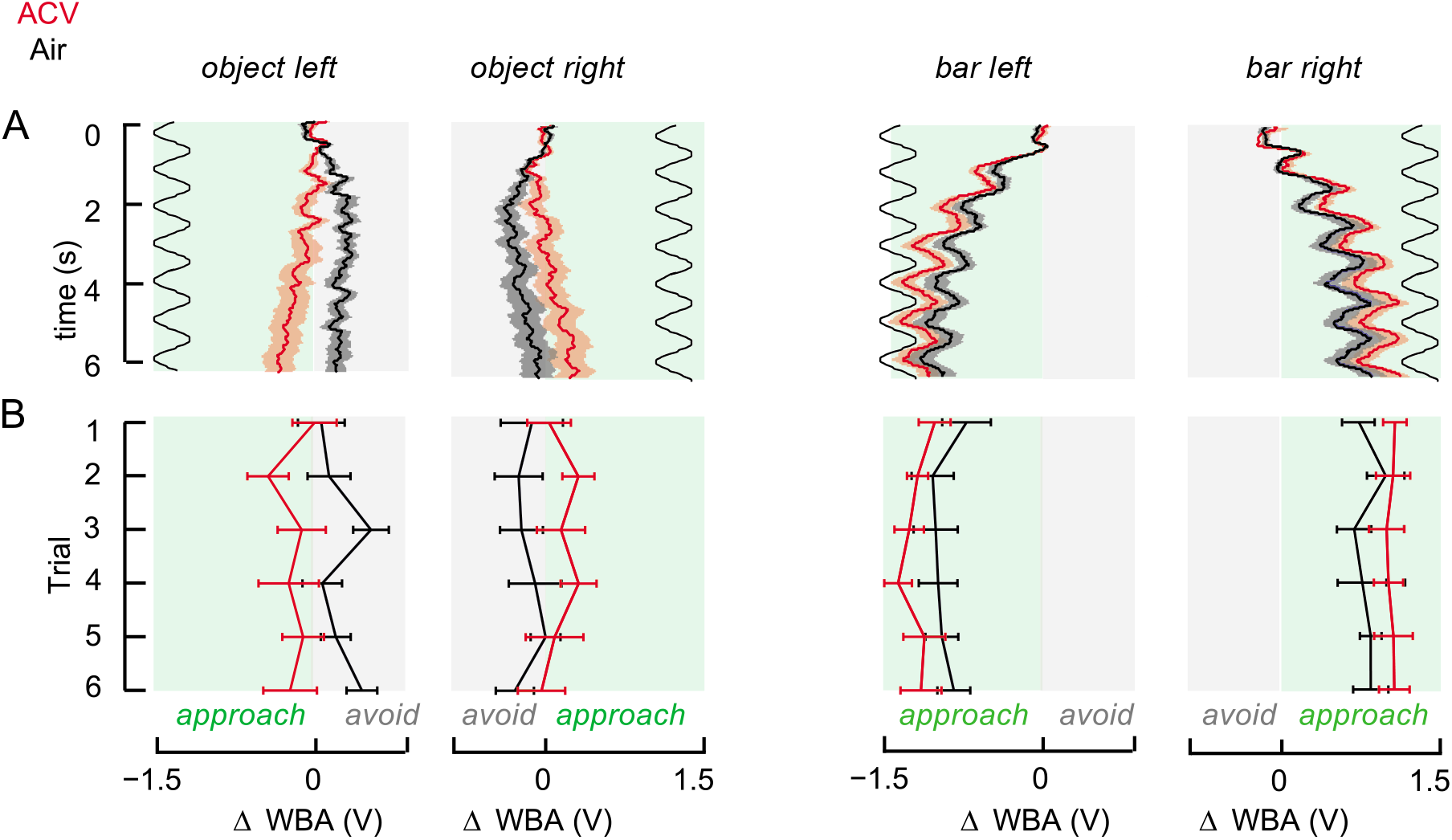
Odor-induced object tracking behavior is observed whether the object is presented on the left or right side of the visual arena. (A) Mean ΔWBA responses (solid lines, n=18) and SEM (shaded regions) of WT-PCF flies to object & bar presented on the left or right side of the visual arena. Red lines indicate responses when the stimulus is presented with ACV, and black lines indicate responses to the visual stimulus in air. Green rectangles denote region of the ΔWBA that correspond to “approach” behavior, which corresponds to the side of the arena that the visual stimulus is presented on. Gray rectangles represent region of the ΔWBA that correspond to “avoid” behavior. Black waveforms represent the 1Hz sine wave that was used to oscillate the visual stimulus ±15° in the left or right front quadrants of the arena. (B) Mean epoch (last 2 seconds) ΔWBA and SEM of single, consecutive trials, averaged across flies (n=18).

**S2.**
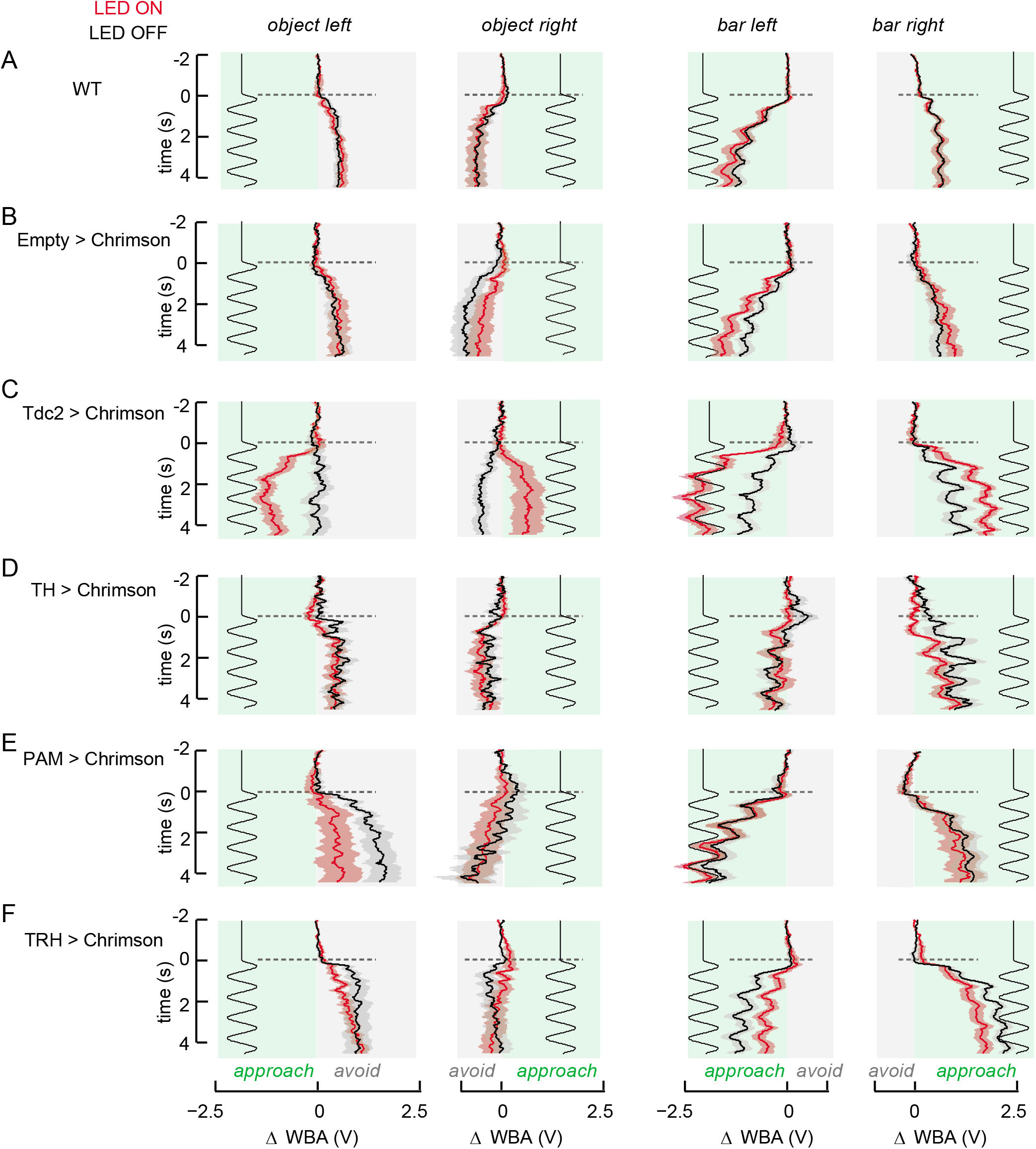
ΔWBA responses of optogenetic-activated aminergic neurons. Mean ΔWBA (solid line) and SEM (shaded regions) for each genotype to each of the four possible stimulus + arena side condition plotted in similar format as Figure S1. Horizontal dashed line represents the onset of visual stimulus motion. Black waveforms represent the 1Hz sine wave that was used to oscillate the visual stimulus ±15° in the left or right front quadrants of the arena. Each panel row shows the responses of one distinct genotype. (A) n=11; (B) n=12; (C) n=16; (D) n=13; (E) n=12; (F) n=16

**S3.**
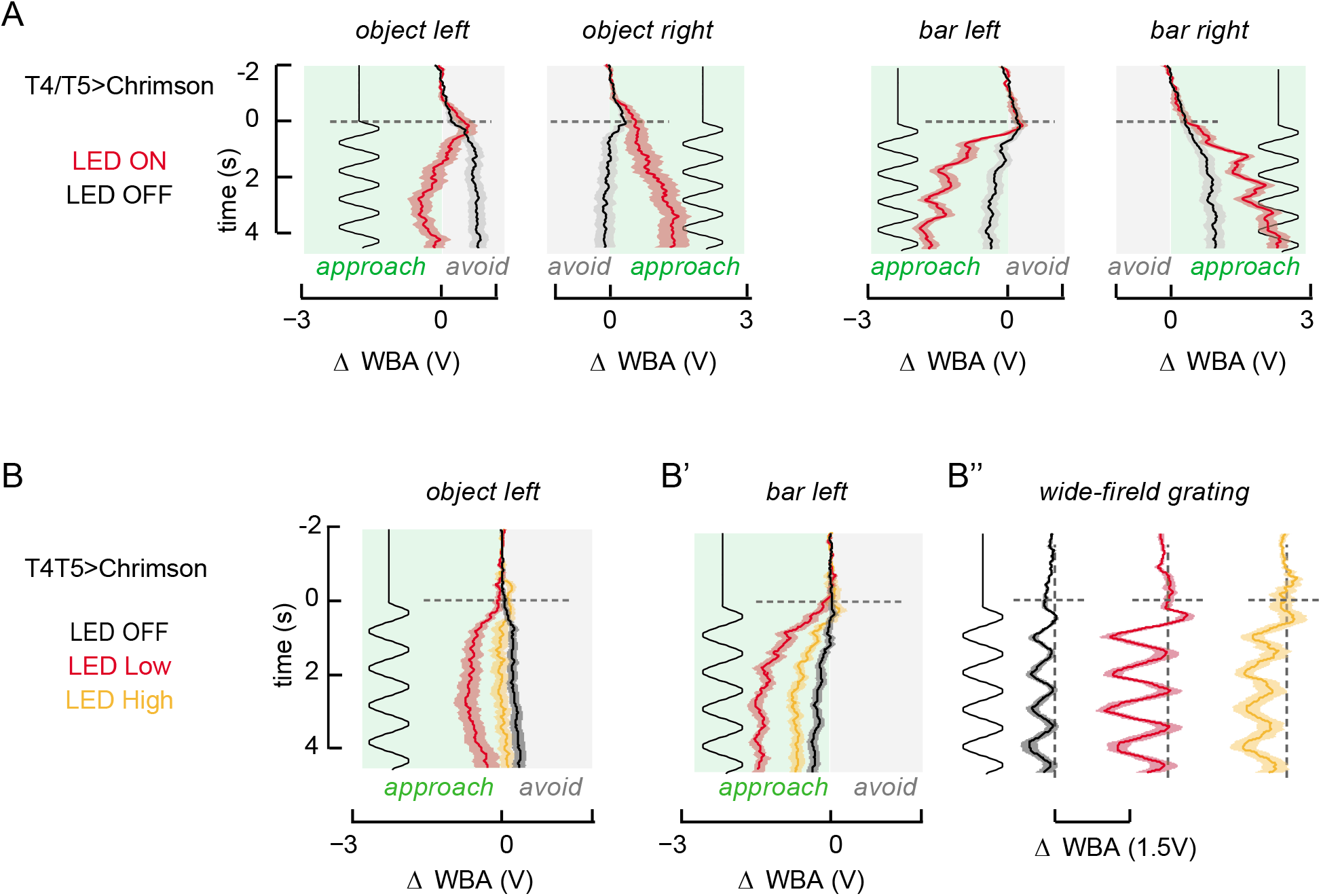
Mean ΔWBA responses of T4/T5 experiments. (A) Mean ΔWBA (solid line) and SEM (shaded regions) for optogenetic activation of T4/T5 (n=18), plotted in the same format as described in Figure S2. (B) The effect of optogenetic activation of T4/T5 neurons (n=12) is dependent upon LED power intensity. High LED intensity (0.040 mW/mm2) abolishes odor-induced object tracking (yellow), while a lower LED intensity (0.010 mW/mm2), which is used in the aminergic optogenetics experiment also, phenocopies odor-induced object tracking (red), similar to Figure 4A. High LED intensity also diminishes bar tracking (B’) and wide-field grating response (B”). These are a separate set of experiments in addition to that presented in Figure 4.

